# The total fungal microbiome functionality

**DOI:** 10.1101/804757

**Authors:** Robert Starke, Petr Capek, Daniel Morais, Nico Jehmlich, Petr Baldrian

## Abstract

Unveiling the relationship between taxonomy and function of the microbiome is crucial to determine its contribution to ecosystem functioning. However, while there is a considerable amount of information on microbial taxonomic diversity, our understanding of its relationship to functional diversity is still scarce. Here we used a meta-analysis of 377 completely annotated and taxonomically different fungal genomes to predict the total fungal microbiome functionality on Earth with all known functions from level 3 of KEGG Orthology using both parametric and non-parametric estimations. The unsaturated model described the accumulation of functions with increasing species richness significantly better (P-value < 2.2e−16) than the saturated model suggesting the presence of widespread and rare functions. Consistent with the parametric approach, the non-parametric Chao-1 estimator that assumes a maximum functional richness did not reach a plateau. Based on previous estimates of fungal species richness on Earth, we propagated the unsaturated model to predict a total fungal microbiome functionality of 42.4 million. Of those, only 0.06% are known today since the vast majority belongs to yet unknown rare functions. Logically, our approach not only highlighted the presence of two types of functions but pointed towards the necessity of novel and more sophisticated methods to unveil the entirety of functions to fully understand the involvement of the fungal microbiome in ecosystem functioning.

---

Ecosystem functioning ^1–4^ is mediated by biochemical transformations performed by a community of microbes from every domain of life ^5^; the microbiome. In every community, multiple organisms from different taxonomic groups can play similar if not identical roles in ecosystem functionality, the so-called functional redundancy ^6^. In fact, functional redundancy of certain functions was shown to be very high with several hundreds to thousands of different taxa expressing the same function within one habitat ^7^. These functions can be statistically inferred based upon homology to experimentally characterized genes and proteins in specific organisms to find orthologs in other organisms present in a given microbiome. This so-called ortholog annotation is performed in KEGG Orthology (KO) ^8,9^ that covers a wide range of functional classes (level 1 of KO) comprising cellular processes, environmental information processing, genetic information processing, human diseases, metabolism, organismal system, *brite* hierarchies and functions not included in the annotation of the two databases *pathway* or *brite*. Logically, however, the bottleneck of describing microbiome functions is the low number of fully annotated specific organisms as they are limited to those that have undergone isolation and extensive characterization while the vast majority of organisms were not yet studied ^10,11^ and the annotation is based on the similarity to the genomes of the very few studied model organisms. As a consequence, fungal microbiome functionality can be inferred based on the composition of the fungal microbiome and its relation to functional parameters ^12^ as indicated by the frequent use of 18S and ITS2 metabarcoding (5,990 publications with the keyword “18S sequencing” and 2,466 with “ITS2 sequencing” in PubMed as of October 3rd 2019). However, as the mere description of the fungal community structure cannot directly assess functionality despite its proficient use to find variables driving the abundance of certain taxa or across multiple taxa down to the species level in complex communities, recently shotgun sequencing of genes (8,857) and transcripts (514) as well as metaproteomics (426) became more and more popular as a direct link between taxonomy and function. However, our understanding of functional diversity and its relationship to taxonomic diversity is still scarce. Here, we use both parametric and non-parametric estimators of functional richness to unveil the relationship between taxonomy and function in fungi with the aim to predict the total fungal microbiome functions on Earth. For this, we downloaded all taxonomically diverse and completely annotated fungal genomes (n=377) from the integrated microbial genomes and microbiomes (IMG) of the Joint Genome Institute (JGI) on August 7^th^ 2019 with taxonomic annotation on species level and functional annotation on level 3 of KO. The parametric estimation comprised of an accumulation curve (AC) ^13^ of increasing functions with increasing species using 1,000 random permutations and its fit to a saturated and an unsaturated model of the Michaelis-Menten kinetics. Otherwise, Chao-1 for every 10% of species richness each with 20 replicates represented the non-parametric estimator. We hypothesized limited functionality with a plateau at high species richness and thus a better fit of the saturated model of the parametric approach and a stagnating Chao-1 estimator with increasing species richness.

The gene count of fungal phyla was not significantly (P-value > 0.5) different than the average gene count of all fungi (**Figure 1a**). Admittedly, the sequenced genomes are heavily in favor of *Ascomycota* (226 = 60.0%) and *Basidiomycota* (122 = 32.4%), and the sequencing of new genomes from the other phyla might reveal significant differences in genome size. Otherwise, significant (P-value < 0.5) differences were found in the number of KO functions were *Ascomycota* (2,666±222 KO functions) comprised of more functions than *Basidiomycota* (2,549±191) that, in turn, had more KO functions compared to *Blastocladiomycota* (2,071±268), *Chytridiomycota* (2,001±276), *Mucoromycota* (2,261±444) and *Zoopagomycota* (2,062±277) (**Figure 1b**). Higher functional diversity in *Ascomycota* and *Basidiomycota* who belong to the subkingdom Dikarya may relate with their classification as higher fungi ^14^ but requires the sequencing and full annotation of additional genomes especially from the other phyla for validation. In the 377 analyzed fungal species, the median of interspecies functional redundancy was found to be 0.03±0.41 (**Figure 1c**). Interspecies redundancy describes the performance of one metabolic function by multiple coexisting and taxonomically distinct organisms ^15^. In fact, most major biogeochemical reactions are driven by a limited set of metabolic pathways that are found in a variety of microbial clades ^16^. Consistent, taxonomic diversity strongly correlated with functional diversity and many ectomycorrhizal fungal species with similar ecological effects co-occurred in the same community ^17^ implying a high interspecies redundancy to mobilize nutrients from organic compounds ^18,19^. Here, interspecies redundancy was either high as 1,592 KO functions were more than 90% of the fungi or low with 4,537 KO functions in less than 10% of the species; together describing 77.3% of all KO functions. Hence, functions appear to diverge into highly redundant across fungal species or unique to only a few. The median of intraspecies functional redundancy was 2.0±1.1 gene copies per KO functions (**Figure 1d**). Generally, low intraspecies redundancy could derive from different KO functions performing functionally similar processes. In fact, all malate dehydrogenases perform the same metabolic function but are annotated by different KO functions (K00024-K00029) due to their involvement in a variety of metabolic pathways. The maximum intraspecies redundancy was found for the ascomycete *Coccidioides immitis* RS that comprised 118 gene copies of the gustatory receptor (K08471) followed by 35 gene copies for the prolyl 4-hydroxylase (K00472, EC 1.14.11.2) and 29 gene copies for the glutathione S-transferase (K00799, EC 2.5.1.18); all of which are functions with low interspecies but high intraspecies redundancy. Noteworthy, the intraspecies redundancy of *C. immitis* RS of 1.6±2.5 was lower compared to the other fungi in the database, which could relate to a rather uncommon lifestyle with a higher share of unique but not essential functions. Indeed, comparative genomic analysis revealed that *C. immitis* is a primary pathogen of immunocompetent mammals ^20^. Functions with high interspecies and high intraspecies redundancy included the yeast amino acid transporter (K16261) with 14.5±8.2 gene copies found in 351 of the 377 fungal species (93.1%), the salicylate hydroxylase (K00480, EC 1.14.13.1) with 13.9±9.5 gene copies (81.7%) and the glutathione S-transferase (K00799, EC 2.5.1.18) with 12.7±11.7 gene copies (98.9%). All of the above belong to the maintenance apparatus of the fungus, namely the transport of amino acids, the incorporation/reduction of oxygen by salicylate hydroxylase, and the detoxification of xenobiotic substrates by glutathione S-transferase. Hence, functions with high interspecies and high intraspecies redundancy are both widespread and essential to every fungus. Functions with high interspecies and low intraspecies redundancy were not found in the 377 genomes. Logically, there might not exist widespread functions that are not essential. Surprisingly and different to our hypothesis, the unsaturated model described the AC significantly better (P-value < 2.2E-16) than the saturated model with both lower Akaike information criterion (AIC) and residual sum of squares (**Table 1**). The unsaturated model is described by the maximum functional richness *f_max_* of 4,716±18*** across the 377 fungal species with an accretion rate *A*_*f*_ of 2.1±0.1*** per fungal species, consistent with the estimate of intraspecies functional redundancy (**Figure 2a**). However, the relationship did not plateau as indicated by the constant of the additive term *k* that is 9.0±0.1***. As it was suggested by the interspecies redundancy, the better fit of the unsaturated model inferred the presence of two types of microbiome functions. On the one hand, widespread functions rapidly increase with the number of species and are ubiquitously abundant in every living fungi. In total, nine functions were found in all of the 377 fungal species. All of which are crucial to sustain life such as the ribose-phosphate pyrophosphokinase (K00948, EC 2.7.6.1) necessary for nucleotide synthesis, the citrate synthase (K01647, EC 2.3.3.1) of the TCA cycle or the superoxide dismutase (K045654, EC 1.15.1.1) that is an important antioxidant defense mechanism. The number of widespread functions is limited and the majority have been identified thus far amounting to, in total, 3,593±31 functions (with 3,534-3,654 as 95% confidence intervals); nearly half of all known functions as of today. Otherwise, roughly 4,300 functions are rare and increase at a much slower rate with an increasing number of species but require time and the evolution of “dead ends”, i.e. species that were unable to evolve a particular function. The addition of more fungal genomes may increase the interspecies redundancy but it is questionable if a function only found in a few of the 377 fungal species can potentially be widespread amongst fungi. Given a species richness of 3.8 million fungi on Earth ^21^ and assuming that the yet unknown fungal microbiome functions are indeed rare, the propagation of the unsaturated model predicted the total fungal microbiome functionality on Earth to be 42,373,186±459,560 (with 41,574,275-43,376,938 as 95% confidence intervals). This estimate was validated by using random subsets of 10, 20, 30, 40, 50, 60, 70, 80 and 90% of all 377 fungal species, which yielded to a plateau of predicted functions when at least 70% of the species were used (**Table 2**). Consistent with the better fit of the unsaturated model from the parametric approach, the non-parametric estimator of functional richness Chao-1 that assumes the existence of a maximum functional richness showed no plateau with increasing species richness of the lower bound estimate (**Figure 2b**) implying that the given species richness is too little to reach a potential maximum of functionality. Admittedly, the exploration beyond the limits of the data is likely to be imprecise and the predictions of fungal microbiome functionality may differ once additional fungal genomes with potentially new functions have been added. However, as of today, our understanding of fungal microbiome functionality is likely limited to a marginal part of all functions.

**Figure 1:**
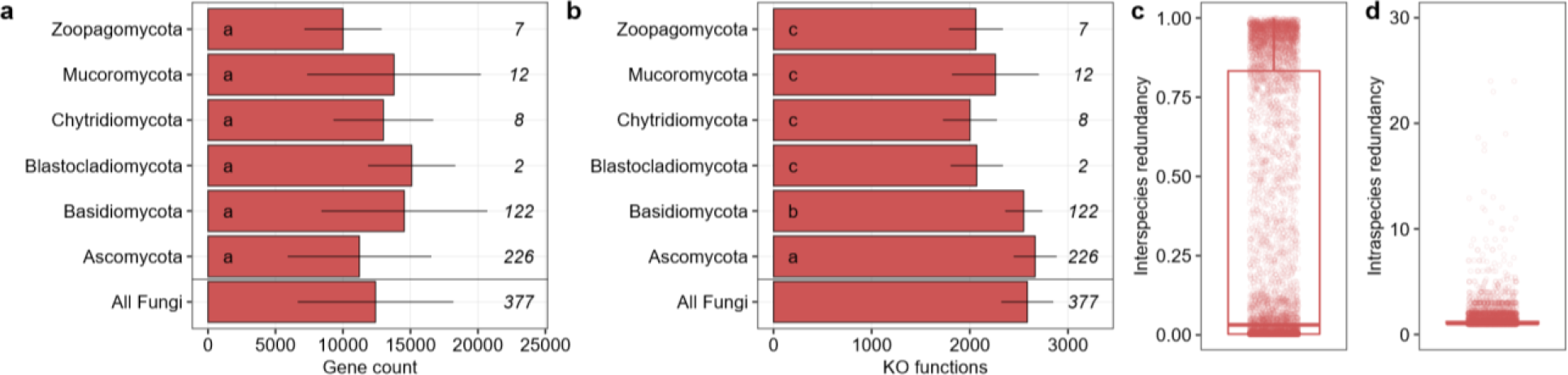
Gene counts (a) and the number of KO functions (b) in fungal phyla given as average with standard deviation. The number of fungal genomes per phylum is given in italic. Data followed by the same letter is not statistically different according to the HSD-test (P-value > 0.5). Total share of KO functions within fungi relative to the total number of fungal species in the database as interspecies redundancy (c) and the number of replicated KO functions within one fungal species in the database as intraspecies redundancy (d).

**Table 1:** The fit of the saturated (S) and the unsaturated model (US) of the accumulation curve indicated by the Akaike’s An Information Criterion (AIC) and residual sum of squares (Res. Sum Sq), the P-value that describes the significant difference between the saturated and the unsaturated model, and the mean prediction with standard deviation (SD) and 95% confidence intervals (CI) at 3.8 million fungal species.

| Model | AIC | Res. Sum Sq | P-value | Prediction | SD | Lower CI | Higher CI |
| --- | --- | --- | --- | --- | --- | --- | --- |
| S | 5,843.751 | 1.17E+08 |  |  |  |  |  |
| US | 4,693.567 | 5514694 | < 2.2E-16 | 42,373,186 | 459,560 | 41,574,275 | 43,376,938 |

**Table 2:** The mean prediction of total fungal microbiome functionality with standard deviation (SD) and 95% confidence intervals (CI) given a richness of 3.8 million fungal species on Earth estimated by the Monte Carlo simulation when random subsamples of the 377 fungal species are used. The prediction stabilizes at around 40 million functions with at least 70% of the fungal species.

| Species (%) | Prediction | SD | Lower CI | Higher CI |
| --- | --- | --- | --- | --- |
| 377 (100) | 42,373,186 | 459,560 | 41,574,275 | 43,376,938 |
| 339 (90) | 37,575,880 | 658,902 | 36,285,040 | 38,868,781 |
| 302 (80) | 46,712,785 | 732,903 | 45,274,956 | 48,152,062 |
| 264 (70) | 38,412,964 | 315,475 | 37,794,156 | 39,031,118 |
| 226 (60) | 62,515,150 | 2,085,421 | 58,421,494 | 66,607,622 |
| 189 (50) | 63,362,037 | 1,645,403 | 60,134,332 | 66,591,118 |
| 151 (40) | 77,516,964 | 1,427,367 | 74,713,805 | 80,321,171 |
| 113 (30) | 97,942,669 | 7,026,524 | 84,131,304 | 111,757,847 |
| 76 (20) | 104,882,008 | 11,188,137 | 82,875,937 | 126,831,401 |
| 38 (10) | 107,435,329 | 3,650,675 | 100,235,159 | 114,636,999 |

**Figure 2:**
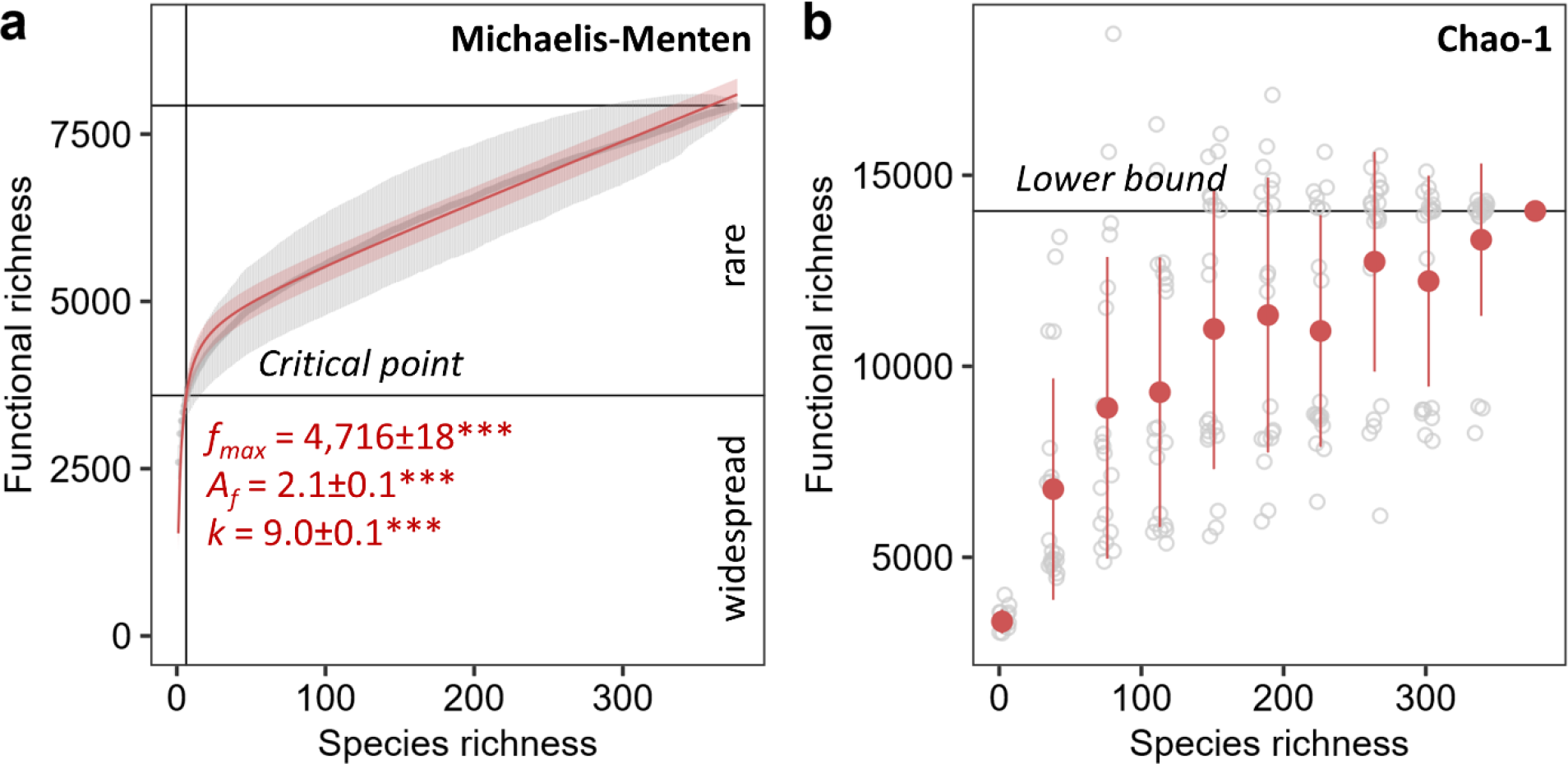
Parametric (a) and non-parametric (b) estimation of total functional richness. The unsaturated model of the accumulation curves as grey points with error bars for the total known fungal microbiome functions derived from the KO database by 1,000 random permutations for every one species richness with 95% confidence intervals. The maximum functional richness is represented by *f*_*max*_, *A*_*f*_ is the accretion rate of functions with increasing number of species, and *k* is the constant of the additive term. Significance of the parameter estimates are indicated by asterisks (*** equals P < 0.001). The Chao-1 was calculated using 20 replicates shown in grey for every 10% of the total fungal species richness in the database starting with two species.

Taken together, we suggest the presence of two types of fungal microbiome functions; widespread and rare functions. Our predictions revealed a potential for million more yet unknown rare functions that, logically, can only be unveiled by novel and more sophisticated methods. However, due to the vast amount of yet unknown functions, it is questionable if the relationship between taxonomy and function is in fact explained by an unsaturated model, if only two types of functions exist and if it is similar when different tools for the functional annotation are used. Noteworthy, the total bacterial microbiome functionality was not predicted as more than 70,000 bacterial genomes must be downloaded individually. At a rate of roughly a minute per genome, the estimated time to gather the information is more than 1,167 hours (or 146 work days). However, given the 100 million bacterial species on Earth ^22,23^ together their lifestyle as micro-environment niche specialists ^24^, the total bacterial microbiome functionality is likely to be much higher and hence, our understanding of the involvement of microbes in ecosystem functioning even lower.

## Materials and Methods

### Metadata collection of the total known fungal microbiome functions

To quantify the relationship between taxonomy and function, all genomes from taxonomically diverse fungal species (as taxonomic unit) were downloaded from the integrated microbial genomes and microbiomes (IMG) of the Joint Genome Institute (JGI) on August 7^th^ 2019, containing the functional annotation from the level 3 of KEGG Orthology ^8,9^ (as functional unit) with counts of every gene per every genome. In total, the database comprised of 377 completely annotated fungal genomes with 7,926 KO functions. The gene counts and KO functions per fungal phylum and in all fungi were retrieved from the database. Interspecies redundancy was calculated as the number of KO functions covered by one randomly chosen species compared to the total number of functions in all species. Intraspecies redundancy or gene redundancy ^25^ was estimated as average of genes per individual KO function in any one species.

### Accumulation curves (AC)

Fungal species were randomly added in intervals of one up to the maximum species richness of 377 with 1,000 random permutations per step using the function *specaccum* from the R package *vegan* ^26^. The AC of the database permutation was then fitted to a saturated (**Equation 1**) and an unsaturated model (**Equation 2**) with the critical point estimated by the term 3*A*_*f*_ as previously described ^27^. The fit of the models was compared by analysis of variance (ANOVA) and Akaike Information Criterion (AIC) ^28^ with a penalty per parameter set to *k* equals two. The total number of KO functions in fungi on Earth was predicted using the global species richness estimate of 3.8 million fungi ^21^ and the Monte Carlo simulation of the function *predictNLS* in the R package *propagate* ^29^. To validate the Michaelis-Menten approach, random subsets of the 377 fungal species with different sizes were used to predict the total microbiome functions as described before. In addition, the non-parametric estimation of the lower bound of functional richness was calculated by Chao-1 again using random subsets of the 377 fungal species with different sizes and 20 replicates each (**Equation 3**).

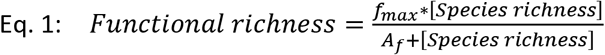

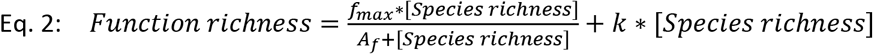

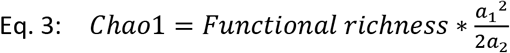

Here, *f*_*max*_ is the maximum functional richness, *A*_*f*_ the accretion rate of functions with an increasing number of species and *k* the constant of the additive term. Functions found only once or twice are indicated by a_1_ as singletons and a_2_ as doubletons, respectively.

## Acknowledgements

RS thanks the Czech Science Foundation for the project 18-25706S. The authors thank Iñaki Odriozola for the determination of the accumulation curves.

## Contributions

RS designed the study. DM and RS performed the computational analysis. PC and RS modelled the data. RS and NJ reviewed the analysis. The paper was written by RS, reviewed by all authors which approved the final version of the manuscript.

